# *CFH* loss in human RPE cells leads to inflammation and complement system dysregulation *via* the NF-κB pathway

**DOI:** 10.1101/2021.04.19.440385

**Authors:** Angela Armento, Tiziana L Schmidt, Inga Sonntag, David Merle, Mohamed Ali Jarboui, Ellen Kilger, Simon J. Clark, Marius Ueffing

## Abstract

Age-related macular degeneration (AMD), the leading cause of vision loss in the elderly, is a degenerative disease of the macula, where retinal pigment epithelium (RPE) cells are damaged in the early stages of the disease and chronic inflammatory processes may be involved. Besides ageing and lifestyle factors as drivers of AMD, a strong genetic association to AMD is found in genes of the complement system, with a single polymorphism in the complement factor H gene (*CFH)*, accounting for the majority of AMD risk. However, the exact mechanism by which *CFH* dysregulation confers such a great risk for AMD and its role in RPE cells homeostasis is unclear. To explore the role of endogenous *CFH* locally in RPE cells, we silenced *CFH* in human hTERT-RPE1 cells. We demonstrate that endogenously expressed *CFH* in RPE cells modulates inflammatory cytokine production and complement regulation, independent of external complement sources or stressors. We show that loss of the factor H protein (FH) results in increased levels of inflammatory mediators (*e*.*g*. IL-6, IL-8, GM-CSF) and altered levels of complement proteins (*e*.*g. C3, CFB* upregulation and *C5* downregulation) that are known to play a role in AMD. Moreover, we identified the NF-κB pathway as the major pathway involved in the regulation of these inflammatory and complement factors. Our findings suggest that in RPE cells, FH and the NF-κB pathway work in synergy to maintain inflammatory and complement balance and in case either one of them is dysregulated, the RPE microenvironment changes towards a pro-inflammatory AMD-like phenotype.

## Introduction

The specific anatomic structure of the human eye permits a tightly regulated local immune response to be sufficient in protecting the retina from external pathogens and to maintain its visual function (1). In immune privileged organs like the eye an excessive immune response, and the subsequent recruitment of circulating immune cells, may lead to tissue damage and affect the function of such a highly specified organ. Ocular immune privilege is mainly ensured by the physical barrier formed by the retinal pigment epithelium (RPE) cell monolayer sitting on an extracellular matrix (ECM), called Bruch’
ss membrane (BrM), which separates the neurosensory retina from the choroid and choriocapillaris (2). The main advantage of an intact RPE/BrM interface is that it provides an effective barrier for the selectivity of molecular diffusion, especially with regard to a possible systemic inflammatory insult (3). Considering that the composition of BrM relies on the deposition of ECM components from both the endothelium cells of the choriocapillaris and the RPE cells, any disruption to RPE cell homeostasis is deleterious for effective barrier maintenance. Moreover, RPE cells exert several other functions needed for retinal health. RPE cells are not only responsible for the phagocytosis and recycling of photoreceptor outer segments (POS), but they also possess antioxidant activity and actively take up nutrients from, and release discard material into, the BrM (4). Although increased signs of inflammation are observed in several retinal degenerative diseases (5), the combination of RPE cell dysfunction, barrier breakdown and subtle, chronic, inflammation is characteristic for the disease age-related macular degeneration (AMD) (6).

AMD is a progressive degenerative disease of the retina, which leads to patients losing their central vision and, in later stages, suffering blindness (7). AMD affects foremost the elderly population and it is estimated that with increasing life expectancy around 300 million people will be affected by 2040 (8). A hallmark of the disease is the presence of deposits, called drusen, within BrM underneath the RPE cells, which not only impair RPE function but also greatly alter the properties of BrM (9). The events that lead to these changes are not yet fully understood, however it is known that AMD is caused by a combination of ageing, genetic predisposition and lifestyle (10-12). The majority of genetic risk lies in the genes of the alternative pathway of the complement system (13), which is an important part of the innate immune system. The canonical role of the complement system is to recognize and mediate the removal of pathogens, debris and dead cells *via* the activation of the complement proteolytic cascade (14). Clearly, tight regulation of complement activation is required to prevent inflammation-induced tissue damage, especially in an immune privileged organ like the eye (15). At the site of complement activation the release of the cleaved complement factors C3a and C5a, called anaphylatoxins, leads to the recruitment and activation of circulating immune cells such as macrophages and leucocytes (16). Additionally, C3a and C5a activate resident immune cells, like microglia and Muller cells, generating a chronic inflammatory environment, which is observed in AMD (17, 18). Complement dysregulation is not only linked to AMD *via* genetic association. Several complement activation products have also been detected in drusen, as well as in the eyes and in the blood of AMD patients (9, 19, 20). One of the most common genetic risks, accounting for 50% of attributable risk for AMD, corresponds to a polymorphism in the complement factor H (*CFH*) gene that consists of a Tyr to His amino acid substitution at position 402 in the pre-processed factor H protein (FH: position 384 in the mature FH protein) (21, 22). The Y402H polymorphism is also present in the alternative splicing product of the *CFH* gene, the protein called factor H-like protein 1 (FHL-1), which is around a third of the size of FH and found to predominate in BrM (23). FH and FHL-1 are negative regulators of the alternative pathway of the complement system and promote the degradation of C3b, a breakdown product of C3 and the central component of the complement activation amplification loop (24). The AMD high-risk genetic variant *CFH* 402H is believed to be involved in AMD pathogenesis in different ways. Indeed, besides the fact that the FH/FHL-1 402H variant has been associated with increased complement activation (25), the same variant also shows reduced binding affinity to ECM components (*e*.*g*. heparan sulphate) (26), oxidized lipids (*e*.*g*. malondialdehyde MDA) (27) and inflammatory mediators (*e*.*g*. C-reactive protein CRP) (28, 29). Most importantly, the function of the FH/FHL-1 proteins may differ depending on their source and location (24, 30). Indeed, in this regard, the endogenous impact of *CFH* proteins in RPE cells has rarely been investigated. In our recent study, we unraveled a non-canonical function of endogenous FH, as the predominant splice form found in RPE cells. By silencing *CFH* in hTERT-RPE1 cells, we showed that FH loss in RPE cells not only modulates the extracellular microenvironment *via* its regulation of C3 levels, but also has an intracellular impact on the antioxidant functions and metabolic homeostasis of RPE cells (31). In the current study, the same model was employed to investigate the endogenous role of FH in the inflammatory response of RPE cells, since RPE cells actively contribute to the maintenance of the immune privileged status of the eye, and not only *via* their barrier function. In particular, we focused on the interactions between FH, inflammation and the nuclear factor kappa-light-chain-enhancer of activated B cells (NF-κB) pathway in RPE cells. The NF-κB pathway is a known key regulator of inflammation and upon canonical regulation of this pathway, the p65 subunit (RelA) of the NF-κB complex is phosphorylated and translocates to the nucleus, where it promotes the transcription of several NF-κB target genes, including inflammatory cytokines, chemokines and also genes involved oxidative stress response (32). Activation of the NF-κB pathway has been associated with several neurodegenerative diseases, including Alzheimer’s and Parkinson’s disease (33), but also in retinal degenerative diseases, such as diabetic retinopathy (34).

Here, we show that RPE cells are immunocompetent with respect to their ability to express and regulate immune-modulatory genes including cytokines and chemokines. FH loss results in an increase of inflammatory cytokines and chemokines in an NF-κB dependent fashion. Moreover, we discovered that FH loss strongly alters the regulation of other complement genes, again in an NF-κB dependent way, thereby creating a dysfunction in complement pathway regulation. As such, the NF-κB pathway emerged as a major signaling pathway controlling immune competence and response in RPE cells.

## Material and Methods

### Cell culture and experimental settings

The human RPE cell line hTERT-RPE1 was obtained from the American Type Culture Collection (ATCC). Cells were maintained in Dulbecco’s modified Eagle’s medium (DMEM; Gibco, Germany) containing 10% fetal calf serum (FCS; Gibco, Germany), penicillin (100 U/ml), streptomycin (100 µg/ml) in a humidified atmosphere containing 5% CO2. Cells were seeded in complete growth medium without phenol red in 12-well plates and allowed to attach overnight. Gene silencing was performed with Viromer Blue reagent according to the manufacturer’s instructions (Lipocalyx, Germany). Culture medium was substituted with fresh medium and siRNA mixture was added dropwise. We employed a mix of three different double strand hairpin interference RNAs specific for either *CFH* or *RELA* and a negative control (Neg), recommended by the provider (IDT technologies, Belgium). In experiments where double silencing was required (siCFH + siRELA), an additional amount of siNeg siRNA was added in the single silencing samples (siNeg, siRELA and siCFH) to keep equal concentrations. Cells were then maintained in serum free medium for the indicated time and where indicated, medium was supplemented with FH (1 µg/ml), C3 (0.1 µg/ml) or C3b (0.1 µg/ml) (CompTech, Texas, USA) for 48 or 144 hours (h).

### RNA extraction, cDNA synthesis and quantitative RT-PCR

At the indicated time points, total RNA was extracted with PureZOL reagent, according to the manufacturer’s instructions (Bio-Rad Laboratories, USA) and cDNA was synthesized *via* reverse-transcription of 1 μg of RNA using M-MLV Reverse Transcriptase (Promega (Wisconsin, USA). cDNA was used to analyse differences in gene expression by qRT-PCR employing iTaq Universal SYBR Green Supermix (Bio-Rad Laboratories, USA) along with gene specific forward and reverse primers (10 μM) listed in Table 1. PCR protocol includes 40 cycles of: 95°C (5 seconds) and 57°C (30 seconds), carried on CFX96 Real-Time System (Bio-Rad Laboratories, USA). Relative mRNA expression of each gene of interest (GOI, Table I) was quantified using 60S acidic ribosomal protein P0 (PRLP0) as the housekeeping control gene.

**Table I.**
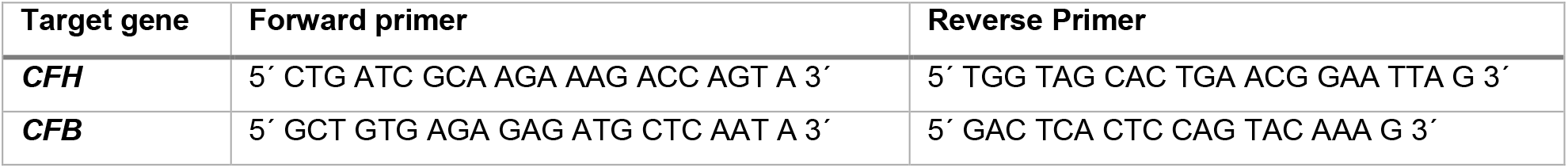
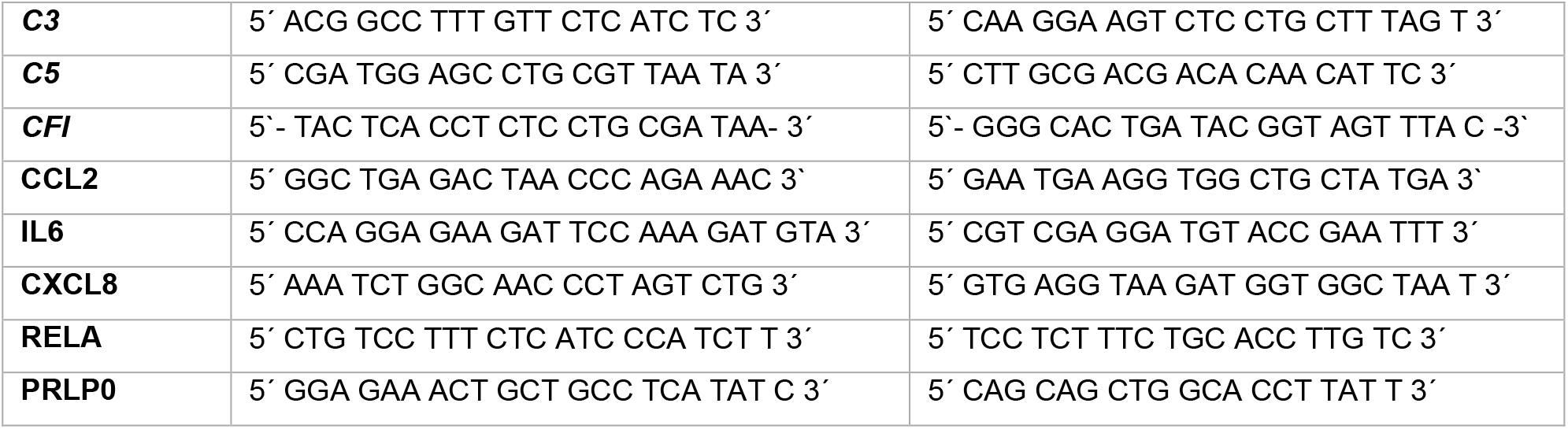
List of primers used in this study.

### Western Blotting

Protein expression was analyzed in both cell lysates and cell supernatants. After debris removal, cell culture supernatants were precipitated with ice-cold acetone. For protein analysis of cell lysates, cells were lysed in Pierce IP Lysis Buffer, containing Halt Protease & Phosphatase Inhibitor (Thermo Fisher, Massachusetts, USA). Protein concentrations were determined with the Bradford quantification assay, using BSA as a standard. Equal amounts of cell lysates or equal volumes of cell supernatants were prepared in NuPAGE LDS Sample Buffer, containing reducing agent (Thermo Fisher, Massachusetts, USA) and analyzed on Novex 8-16% Tris-Glycine gels (Invitrogen, California, USA). Subsequently proteins were transferred onto PVFD membranes and western blot detection carried out as previously described (31), using the primary antibodies listed in Table II. Pictures were acquired with a FusionFX imaging system (Vilber Lourmat, France) and the intensity density of individual bands was quantified using the ImageJ software.

**Table II.** List of primary antibodies used in this study.

| Antibody | Supplier | Number |
| --- | --- | --- |
| **β-actin** | Cell Signaling | #3700 |
| **Complement C3** | Invitrogen, ThermoFisher | #PA5-21349 |
| **Factor H (FH)** | SantaCruz Biotechnology | sc-166608 |
| **p-NF-kB p65** | Cell Signaling | #3033 |
| **Total NF-kB p65** | Cell Signaling | #8242 |

### C3b ELISA

C3/C3b ELISA to evaluate the concentration of C3/C3b in cell culture supernatants was performed according to the manufacturer’
ss instructions (Abcam, UK). Samples and standard controls were loaded in 96 well-plates coated with specific C3b antibody. Absorbance was read at a wavelength of 450 nm immediately after the assay procedure using a Spark multimode microplate reader (Tecan, Switzerland). Subtraction readings at 570 nm were taken to correct optical imperfections.

### Cytokine array

The Proteome Profiler Human Cytokine Array Kit (R&D Systems, Minnesota, USA) was employed to determine the relative levels of 36 different cytokines and the assay was performed according to the manufacturer’s instructions. Briefly, the membranes were blocked for 1 h at room temperature. The cellular supernatant samples were further prepared by mixing 400 μl of the sample with 500 μl Array Buffer 4, 600 μl Array Buffer 5 and 15 μl of Detection Antibody Cocktail and incubated for 1 h at room temperature. The prepared mixture was added to the membranes, followed by an incubation period overnight at 4°C. The membranes were washed 3 x 10 minutes, incubated in diluted HRP-Streptavidin (1:2000) for 30 minutes, and washed again for 3 x 10 minutes. The Chemi Reagent Mix (1:1 ratio) was dropped onto the membranes, incubated for 1 minute, and the signal was detected by FusionFX (Vilber Lourmat, France) in the automatic mode and, additionally, in an individual programmed mode with an increasing detection time of: 0.5, 1, 1.5, 2, 4, 6, 10 minutes. The results were evaluated with the Fusion software and ImageJ by measuring the intensity density.

### Bioinformatic analyses

Data analysis of the raw values from the cytokine array as obtained and measured using ImageJ, were normalized to the positive control. For principal component analysis (PCA), values were Pareto scaled by dividing each variable by the square root of the standard deviation to minimize the effect of small noisy variables.

The Variable Importance in Projection (VIP) in a Partial Least Square Discriminant Analysis (PLS-DA) was used to identify the most discriminative cytokines for each biological group following siRNA treatment. Similar to the PCA analysis and normalized to the positive control, Pareto scaled values were used. PCA and VIP score analysis were carried out using the R package MetaboAnalystR, integrated in the publicly available platform for statistical analysis metaboanalyst.ca (35).

## Statistical analysis

The data are presented as mean with the standard error of the mean (SEM) and were generated and tested for their significance with GraphPad Prism 8 software. All data sets were tested for normal distribution, assessed with Shapiro-Wilk normality test. Depending on normal distribution and the parameters to be compared the following tests were performed: Mann-Whitney test was used in case of non-normal distribution, unpaired student’
ss t-test was used to compare siNeg vs either siCFH or siRELA condition and to compare siCFH vs siCFH + siRELA. Ratio paired t-test was used to compare the relative changes between siCFH vs siCFH+siRELA and siCFH vs siCFH treated (FH, C3 or C3b), only when both conditions were normalized to siNeg control. Values were considered significant with p<0.05.

## Results

### *CFH* loss leads to upregulation of C3 and inflammatory cytokines in RPE cells

In our previous study, we used siRNA silencing of the *CFH* gene to investigate the impact of reduced FH levels and activity in hTERT-RPE1 cells in response to oxidative stress after 48h *in vitro* (31). Here, we used the same model to investigate immune reactivity of RPE cells after *CFH* knockdown and prolonged its silencing period up to 6 days (144h). In this way, we were able to mimic features of an early stage of AMD where the barrier function of RPE cells remains intact, and therefore the protein composition of the retinal microenvironment depends mostly on RPE cells protein production. Using both the short (48h) and the prolonged (144h) silencing period, we investigated the impact of endogenous FH loss on pro-inflammatory cytokine production and on complement system regulation.

Efficient *CFH* silencing after 48h was already shown before and reproduced in this study (31). Here, we show that also after 144h *CFH* mRNA was significantly reduced in *CFH* knock-down (siCFH) RPE cells compared to control cells (siNeg). The silencing efficiency of almost 90% after 48h was maintained also at 144h (Fig. 1a). Likewise, FH protein levels remained almost undetectable in cell culture supernatants of siCFH cells after 144h (Fig. 1b). With prolonged FH loss, *C3* gene expression (Fig. 1e) and C3 protein levels (Fig. 1c-d) increased significantly. Levels of *C3* mRNA were found to be 20-fold higher in siCFH cells at both time points (Fig. 1e). Similarly, protein levels of secreted C3 were significantly elevated in siCFH cells, as shown by a 2-fold increase in C3/C3b ELISA (Fig. 1c) and a 4-fold increase in C3/C3b alpha and beta chains in Western Blot (Fig. 1d).

**Figure 1.**
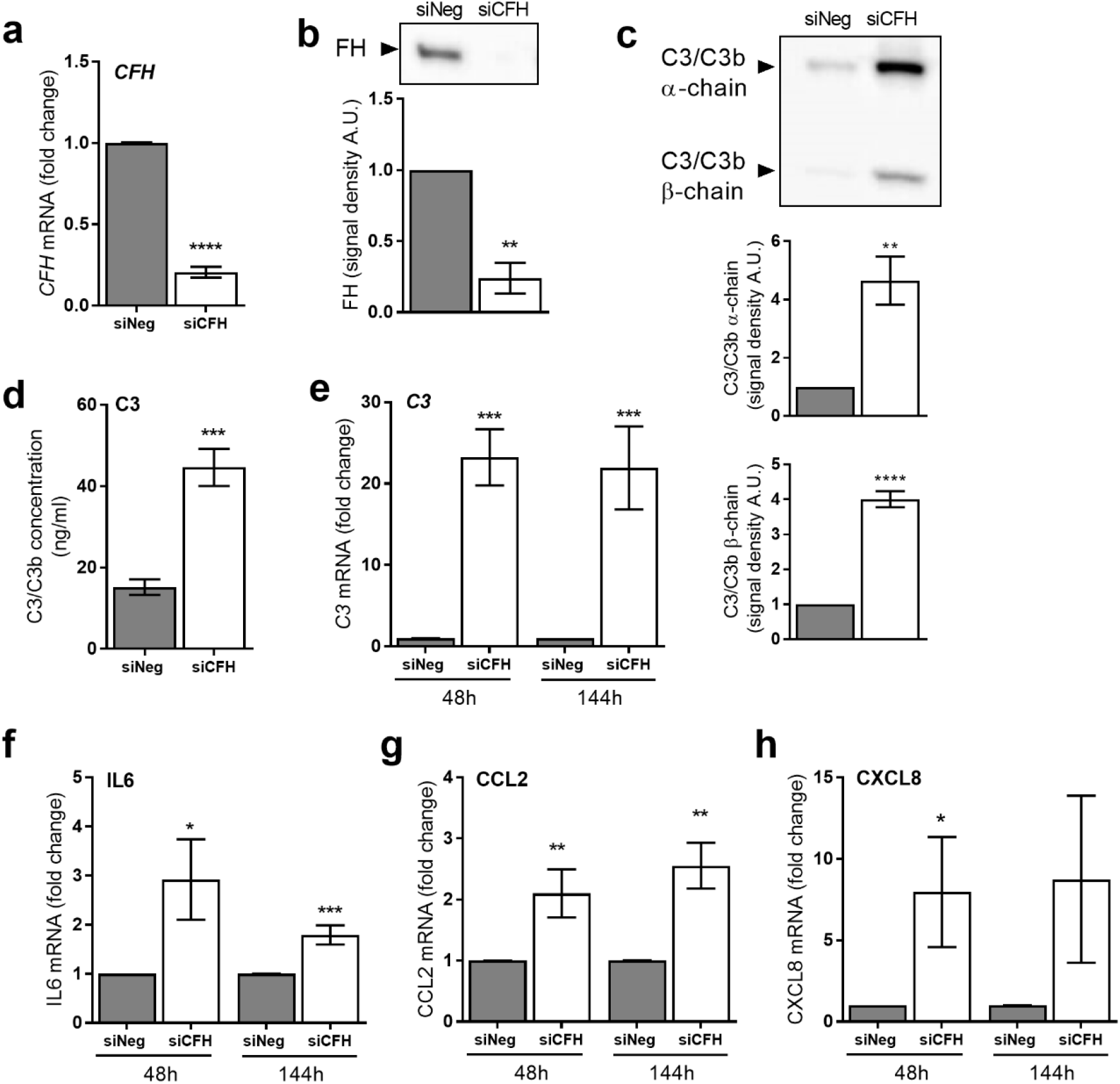
Short and sustained FH reduction leads to increased expression levels of C3 and inflammatory cytokines. hTERT-RPE1 cells were seeded, left to attach overnight and silenced for 24 hours with negative control (siNeg) or *CFH* specific (si*CFH*) siRNA. Afterwards, cells were kept in serum-free medium (SFM) and cell pellets and cell culture supernatants were collected after 48 and 144 hours. **a** Evaluation of *CFH* expression after 144h by qRT-PCR analyses. Data are normalized to the housekeeping gene PRLP0 using Δ ΔCt methods. SEM is shown, n=5. **b** Western blot analyses of FH protein levels in cell culture supernatants of hTERT-RPE1 cells after 144h. Quantification of signal density of 3 independent experiments is shown. **c** Western blot analyses of C3/C3b α-chain and β-chain protein levels in cell culture supernatants of hTERT-RPE1 cells after 144h. Quantification of signal density of 5 independent experiments is shown. **d** C3/C3b ELISA analyses of cell culture supernatants of hTERT-RPE1 cells after 144h. SEM is shown, n=6. **e-h** Monitoring of gene expression by qRT-PCR analyses in hTERT-RPE1 cells: **e** complement component 3 (*C3*), **f** interleukin-6 (IL6), **g** C-C Motif Chemokine Ligand 2 (CCL2) and **h** interleukin-8 (CXCL8). Data are normalized to housekeeping gene PRPL0 using Δ ΔCt method. SEM is shown, n=5-8. Western Blot images were cropped, and full-length blots are presented in Supplementary Fig. S1. Significance was assessed by Student’s t-test. *p<0.05, **p<0.01, *** p<0.001, ****p<0.0001.

Given that early AMD is hallmarked by persistent inflammation (36, 37), we investigated the levels of relevant inflammatory cytokines, including: interleukin-6 (IL6), C-C Motif Chemokine Ligand 2 (CCL2) and interleukin-8 (CXCL8). When FH was downregulated in siCFH cells, we observed an upregulation of IL6 (3-fold at 48h and 2-fold at 144h) and CCL2 (2-fold at 48 and 2.5-fold at 144h) (Fig. 1f-g). Moreover, CXCL8 levels were 8-fold upregulated in siCFH cells, significantly after 48h (Fig. 1h). This indicates that reduction of FH levels and activity in RPE cells leads to an upregulation of relevant pro-inflammatory molecules.

### Cytokine expression mediated by FH loss is driven by NF-κB activity

Changes in RPE gene expression after FH reduction, suggests that a pro-inflammatory pathway may be regulated by FH in RPE cells. As NF-κB plays a major role in regulating a variety of cytokine expression levels (38), we next investigated if the activity of NF-κB was changed by *CFH* silencing. To do so, we monitored the levels of the activated phosphorylated form of the NF-κB p65 subunit and its total levels in siNeg and siCFH RPE cells over time (Fig. 2a). We found a dramatic and sustained increase in the activation levels of NF-κB, as shown by the increased relative ratio of phosphorylated/total NF-κB p65 subunit (Fig. 2b).

**Figure 2.**
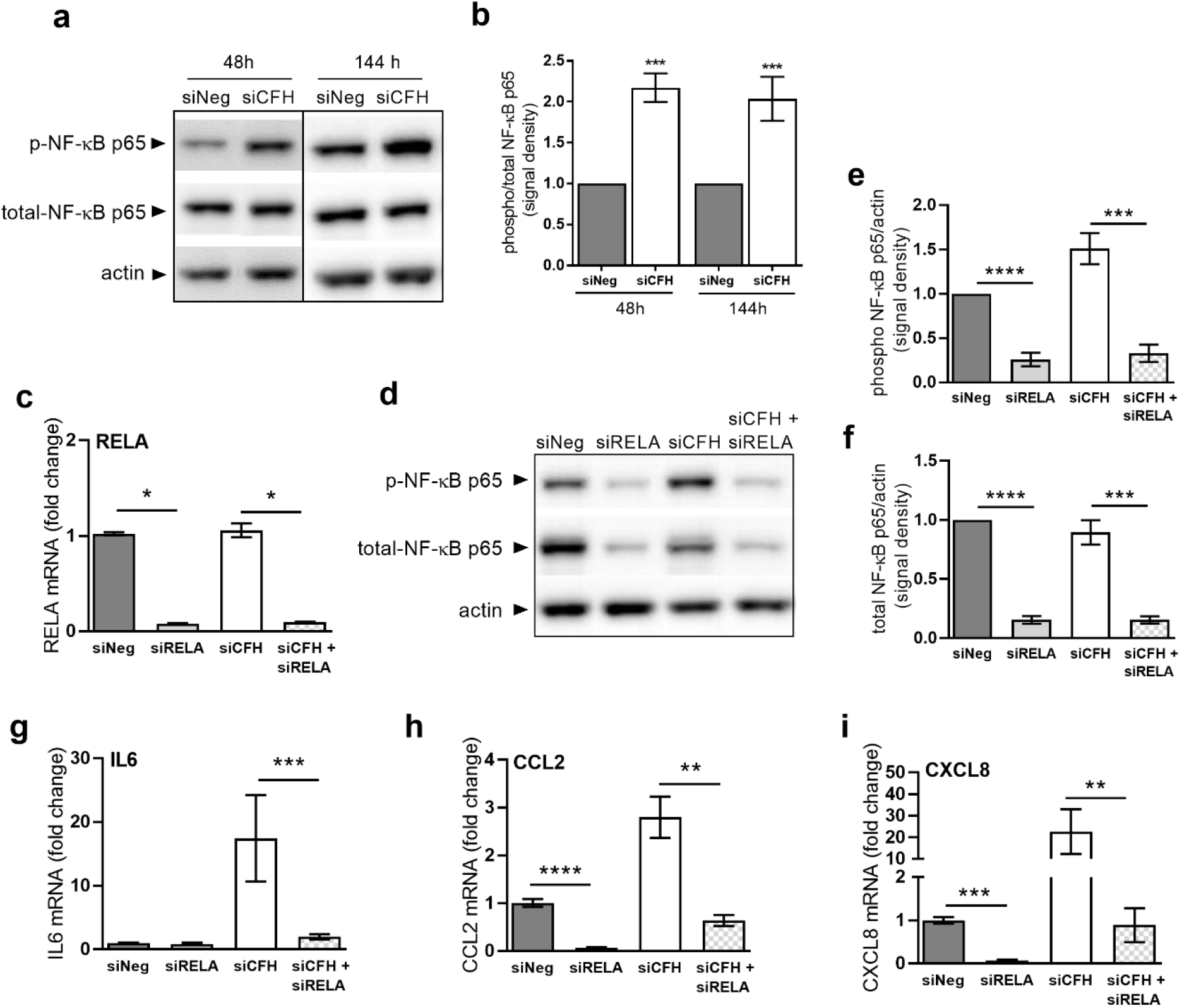
RPE cells deprived of FH show NF-κB activation and blocking NF-κB abolishes the effects of FH loss on cytokine gene expression levels. hTERT-RPE1 cells were seeded, left to attach overnight and silenced for 24 hours with either negative control (siNeg), *CFH* specific (si*CFH*) or NFκB/*RELA* specific (siRELA) siRNA or with a combination of siCFH and siRELA siRNA. Afterwards, cells were kept in serum-free medium (SFM) and cell pellets and cell culture supernatants were collected after 48 and/or 144 hours. **a** Western blot analyses of phosphorylated and total levels of p65 NF-κB subunit in cell lysates of hTERT-RPE1 cells after 48h and144h. Total actin was used as loading control. **b** Quantification of signal density of at least 4 independent experiments as reported in a. Bars indicate the signal density ratio between levels of phosphorylated and total p65 NFκB subunit. **c** Evaluation of RELA gene expression levels by qRT-PCR analyses in hTERT-RPE1 cells after 48h. Data are normalized to the housekeeping gene PRPL0 using Δ ΔCt methods. SEM is shown, n=5. **d** Western blot analyses of phosphorylated and total levels of p65 NFκB subunit in cell lysates of hTERT-RPE1 cells after 48h. Total actin was used as loading control. **e** Quantification of signal density of 3 independent experiments in the conditions reported in c-d. Bars indicate the signal density ratio between phosphorylated p65 NFκB subunit and actin. **f** Quantification of signal density of 3 independent experiments in the conditions reported in c-d. Bars indicate the signal density ratio between total p65 NFκB subunit and actin. **g-i** Gene expression analyses by qRT-PCR of hTERT-RPE1 cells in the conditions reported in c-d: **g** interleukin-6 (IL6), **h** C-C Motif Chemokine Ligand 2 (CCL2) and **i** interleukin-8 (CXCL8). Data are normalized to housekeeping gene PRLP0 using Δ ΔCt method. SEM is shown, n=3-5. Western Blot images were cropped, and full-length blots are presented in Supplementary Fig. S2-3. Significance was assessed by Student’s t-test. *p<0.05, **p<0.01, *** p<0.001, ****p<0.0001.

To investigate a potential direct correlation between reduced FH activity and the observed NF-κB pathway activation, we employed concomitant double silencing of *CFH* and RELA, the gene coding for the NF-κB p65 subunit. First, RELA silencing efficiently reduced the levels of the gene by about 90% in siRELA and siCFH + siRELA cells (Fig. 2c). *CFH* silencing had no impact on the expression level of RELA (Fig. 2c). In RELA silenced cells, the protein levels of NF-κB p65 were also greatly reduced (Fig. 2d). Quantification of protein levels for both the phosphorylated form of NF-κB p65 (Fig. 2e) and total NF-κB p65 (Fig. 2f) in siRELA and siCFH + siRELA cells show a significant reduction of protein abundance.

Next, we evaluated gene expression levels of the identified upregulated cytokines in response to RELA silencing. Interestingly, we found that under control conditions (*i*.*e*. in the presence of FH), the NF-κB pathway regulates the expression of CCL2 and CXCL8, but not that of IL6 (Fig. 2g-i). As shown in Fig. 2g, IL6 levels only rise in the absence of FH activity (siCFH). Conversely, a downregulation of NF-κB p65 in siCFH cells (siCFH + siRELA), lowers the gene expression of all three cytokines back to basal levels (Fig. 2g-i). In particular, a strong significant reduction was observed for IL6 (Fig. 2g), CXCL8 (Fig. 2i) and for CCL2 (Fig. 2h).

Next, we tested whether exogenous application of FH could revert the effects of endogenous siRNA-based *CFH* suppression on both NF-κB activation as well as the expression of inflammatory cytokines. At the same time, we evaluated the effects of an addition of C3 and C3b. The addition of exogenous complement factors, however, did not change NF-κB activation levels (Suppl. Fig. S4a-b) nor gene expression levels of IL6 (Suppl. Fig. S4c) and CCL2 (Suppl. Fig. S4d) at any of the time points tested (48h and 144h).

In order to assess whether the changes in gene transcription would translate into an inflammatory microenvironment outside of the RPE cells, we monitored the levels of secreted inflammatory factors *via* a cytokine array analyzing the serum free conditioned medium supernatant of hTERT-RPE1 cells after 48 hours (Fig. 3a).

**Figure 3.**
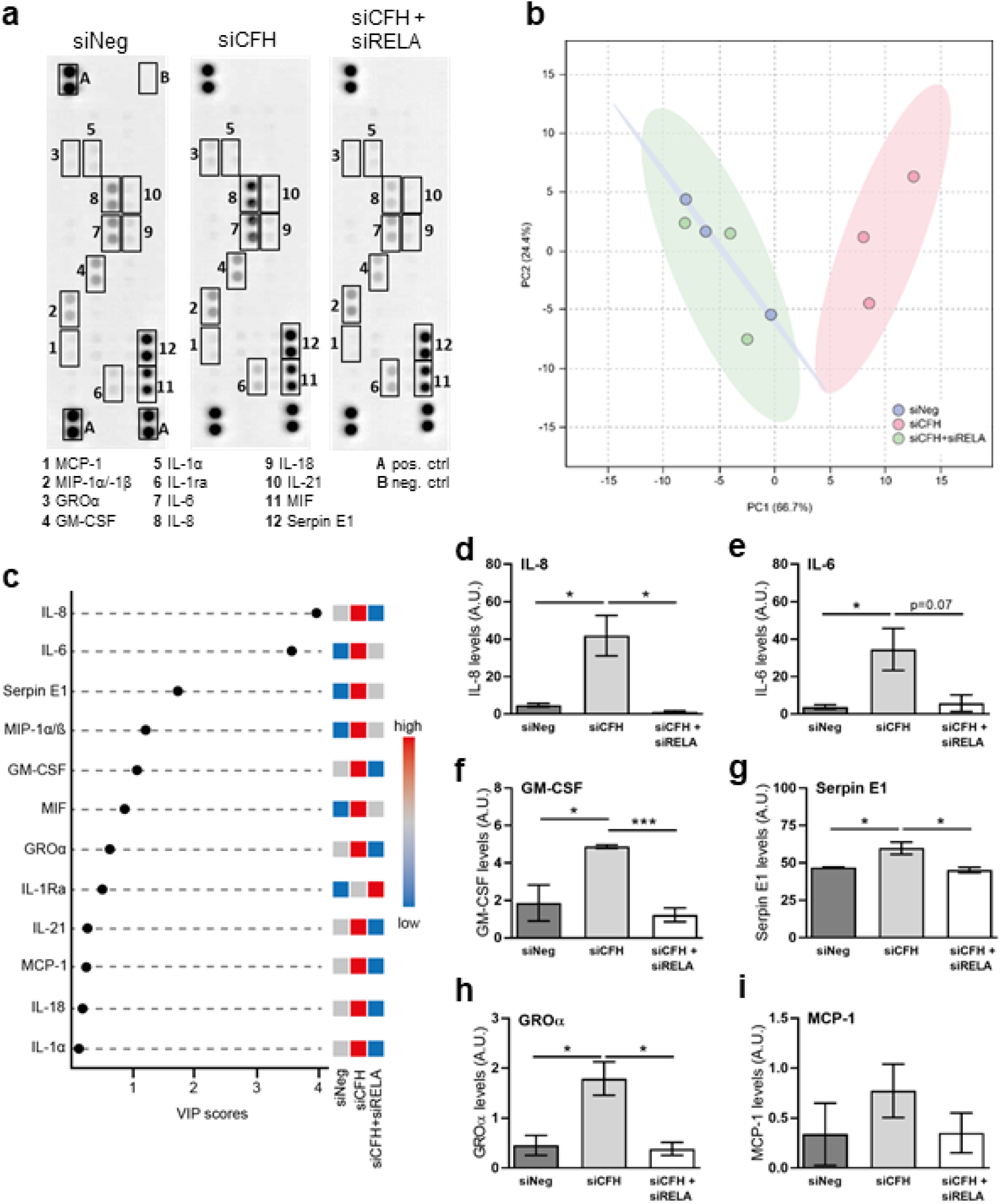
Blockade of NFκB abolishes the effects of FH loss on secreted cytokines. hTERT-RPE1 cells were seeded, left to attach overnight and silenced for 24 hours with either negative control (siNeg), *CFH* specific (si*CFH*) or with a combination of siCFH and NFκB/RELA specific (siRELA) siRNA. Afterwards, cells were kept in serum-free medium (SFM) and cell culture supernatants were collected after 48h. **a** Representative image of a Proteome Profiler Human Cytokine Array analysis performed on cell culture supernatants collected from hTERT-RPE1 cells after 48h. **b** PCA Analysis of the Cytokine array data for all biological replicates, samples were colored according to the corresponding siRNA treatment group, 95% confidence regions were plotted and colored according to each group. **c** Variable importance in projection (VIP) score plot derived from PLS-DA analysis, the top cytokines that contribute to the segregation between the groups were plotted and their differential abundance was color scaled according to their enrichment (red), depletion (blue), or unchanged (grey). **d-i** Quantification of signal density in the conditions reported in a: **d** interleukin-8, IL-8; **e** interleukin-6, IL-6; **f** colony stimulating factor 2, GM-CSF; **g** serpin family E member 1, Serpin E1; **h** C-X-C Motif Chemokine Ligand 1, CXCL1/GROα; **i** C-C Motif Chemokine Ligand 2,CCL2. SEM is shown. n=3. Significance was assessed by Student’s t-test. *p<0.05, **p<0.01, *** p<0.001).

Using PCA analysis, we plotted all the signal intensities for each cytokine from all the biological replicates of each group: siNeg, siCFH and siCFH+siRELA. As shown in Fig. 3b, there is clear segregation between siNeg and siCFH as they cluster apart according to the first principal component (PC1), thus indicating a clear effect of siCFH silencing on the RPE cytokine signature. The combination of siCFH + siRELA had little effect on the cytokine profile signature as siCFH + siRELA group cluster tightly with siNeg control (Fig. 3b). This suggests that downregulation of NF-κB p65 (siRELA) in siCFH silenced RPE cells results in a reversion of the pro-inflammatory phenotype to normal para-inflammatory homeostasis. As the microarray used for analysis only covers a limited number of cytokines and their relative quantification was based on differences in signal detection after blotting, we chose to use a more supervised approach. Variable Importance in Projection (VIP) values from Partial Least Square Discriminant Analysis (PLS-DA) was used to gain a quantitative estimation of the discriminatory power of each individual cytokine (Fig. 3c). VIP score analysis detected 12 cytokines that significantly differentiated between the 3 siRNA groups. IL-8 and IL-6, also shown in Fig. 3d and Fig 3e are the 2 cytokines that contribute the most to the segregation between siNeg, siCFH and siCFH + siRELA. Besides these two, most of the cytokines analysed on the array were increased in the siCFH group when compared to the siNeg controls: colony stimulating factor 2 (GM-CSF, Fig. 3f), serpin family E member 1 (Serpin E1, Fig. 3g), C-X-C Motif Chemokine Ligand 1 (CXCL1/GROα, Fig. 3h), C-C motif chemokine ligand 3 and 4 (MIP-1α/-1β, Suppl. Fig. S6a), while the effects on MCP-1, IL18 and IL-1a were less pronounced and only slightly changed (Fig. 3i). Most of these cytokines exhibit a decreased level of abundance when silencing of FH (siCFH) and NF-κB p65 (siRELA) was combined: IL-8 (Fig. 3d), IL-6 (Fig. 3e), GM-CSF (Fig. 3f), Serpin E1 (Fig. 3g), CXCL1/GROα were reduced to a base level indicating that an inhibition of the NF-κB pathway can dampen or abrogate the consequences of FH loss (Fig. 3h).

An exception to this pattern was seen with interleukin 1 receptor antagonist (IL-1Ra, (Fig 3a, c, and Suppl. Fig. S6b), which antagonizes the inflammatory effects of IL-1α/-1β *via* competitive binding to their receptors. The upregulation of IL-1Ra as an anti-inflammatory cytokine suggests that its expression is negatively regulated by NF-κB pathway activity. Minimal differences were observed in between the conditions (Suppl. Fig. S6) for macrophage migration inhibitory factor (MIF), interleukin-1 alpha (IL-1a), and interleukin-18 (IL-18) and interleukin-21 (IL-21), although the latter was slightly reduced in response to siRELA.

### FH loss alters transcription of complement genes *via* the NF-κB pathway

The transcriptional regulation of complement genes has remained poorly understood in RPE cells, as well as in general. After finding that reduction of FH activity resulted in a marked upregulation of C3 expression (Fig. 1), we investigated whether either suppression of FH expression or NF-κB activity would regulate the expression levels of additional complement factors or regulators.

We observed a significant reduction of *CFH* mRNA levels in siRELA cells (Fig. 4a), followed by reduced levels of FH at the protein level (Fig. 4b). C3 levels were also reduced in siRELA cells, both at the protein level (Fig. 4c) as well as RNA level (Fig. 4d). Addition of exogenous FH could only partially revert the effects of endogenous *CFH* silencing on *C3* mRNA levels (Suppl. Fig. S8a) and C3 secreted levels (Suppl. Fig. S8b) after 144h of incubation. Supplementation with C3 and C3b had no impact (Suppl. Fig. 8a) at any time point.

**Figure 4.**
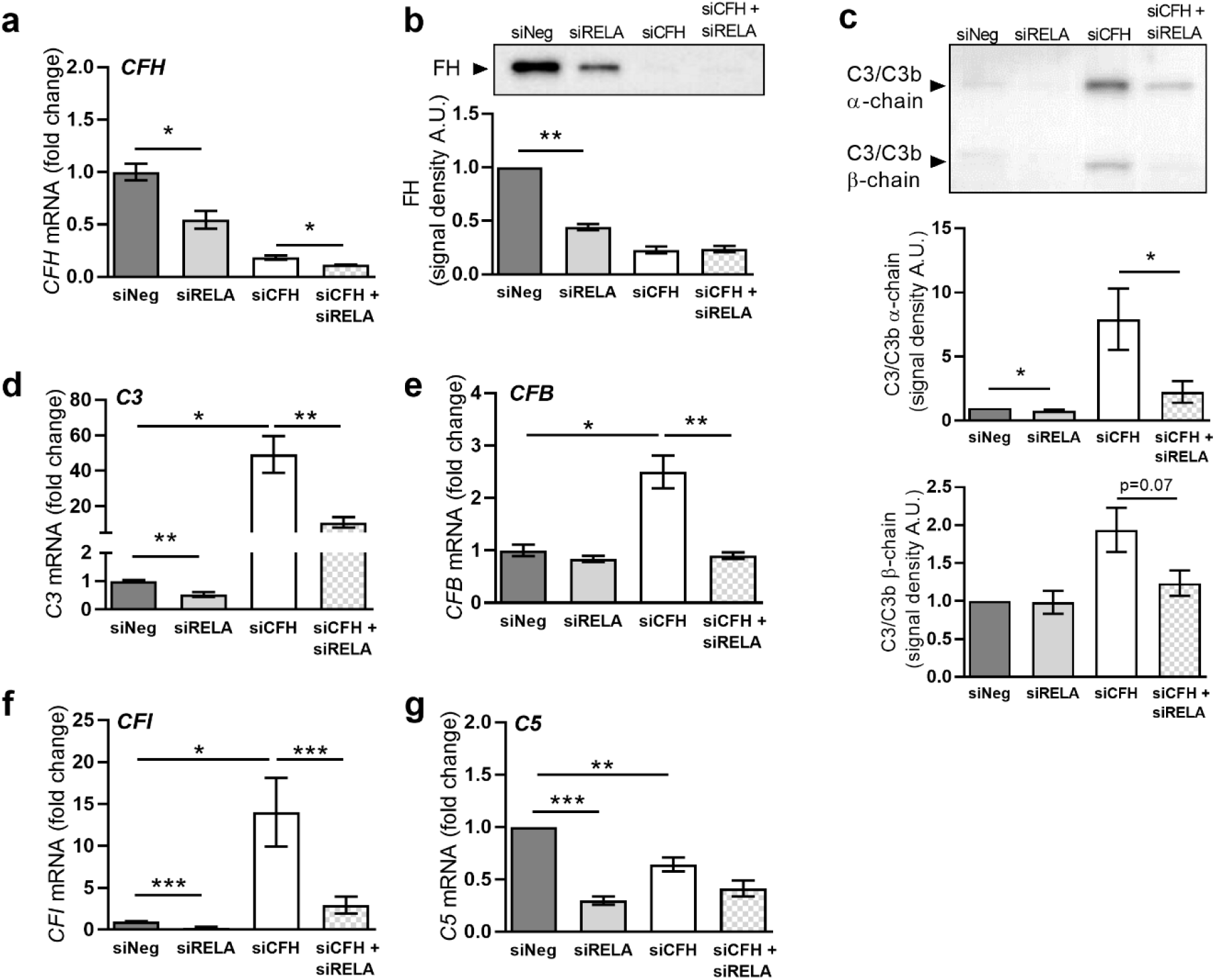
FH loss alters gene transcription of complement system genes *via* the NFκB pathway. hTERT-RPE1 cells were seeded, left to attach overnight and silenced for 24 hours with either negative control (siNeg), *CFH* specific (si*CFH*), NFκB/RELA specific (siRELA) siRNA or with a combination of siCFH and siRELA siRNA. Afterwards, cells were kept in serum-free medium (SFM) and cell pellets and cell culture supernatants were collected for after 48h. **a** Evaluation of *CFH* expression by qRT-PCR analyses. Data are normalized to the housekeeping gene PRPL0 using Δ ΔCt methods. SEM is shown, n=5. **b** Evaluation of *C3* expression by qRT-PCR analyses. Data are normalized to the housekeeping gene PRPL0 using Δ ΔCt methods. SEM is shown, n=5. **c** Western blot analyses of C3 α-chain and β-chain protein levels in cell culture supernatants. Quantification of signal density of 4 independent experiments is shown. **d** Western blot analyses of FH protein levels in cell culture supernatants. Quantification of signal density of 3 independent experiments is shown. **e** Evaluation of *CFB* expression by qRT-PCR analyses. Data are normalized to the housekeeping gene PRPL0 using Δ ΔCt methods. SEM is shown, n=5. Western Blot images were cropped, and full-length blots are presented in Supplementary Fig. S7. Significance was assessed by Student’s t-test. *p<0.05, **p<0.01.

Subsequently, we investigated the gene expression levels of *CFB*, an important positive regulator of the alternative pathway of the complement system. We found a significant 2.5-fold increase in the *CFB* gene after *CFH* silencing and a return to basal levels after silencing NF-κB p65, while *CFB* levels were not affected by siRELA alone (Fig. 4e). Addition of exogenous FH could only partially revert the effects of endogenous *CFH* silencing on *CFB* mRNA levels, while addition of C3 and C3b had no effects (Suppl. Fig. S8c). Next, we evaluated the expression of *CFI*, a negative regulator of the complement activation in the presence of FH as a co-factor. The RNA levels of *CFI* were 12-fold higher in siCFH cells and were significantly reduced in siRELA cells. (Fig. 4f). Furthermore, we analysed the expression of *C5*, another major component of the complement cascade. The levels of *C5* RNA (Fig. 4g) were reduced by both silencing FH expression in siCFH cells as well as after NF-κB p65 silencing in siRELA cells.

Most importantly, all factors belonging to the complement system, whose levels were increased in the absence of FH, were reduced *via* suppression of NF-κB activity. Thus, *CFB* RNA levels were redirected to basal levels (Fig. 4e); as well as a clear reduction of *CFI* RNA levels was observed (Fig. 4f). *C3* RNA levels were significantly reduced by half (Fig. 4b) and C3 secreted protein levels were also significantly reduced (Fig. 4c).

## Discussion

Given the strong association of the *CFH* gene with AMD, and a clear role of RPE cells in maintaining homeostasis in the retinal microenvironment, we investigated the role of FH in RPE cells with respect to its impact on balancing molecular mechanisms of inflammation. Here, we demonstrate that endogenously expressed *CFH* in RPE cells modulates inflammatory cytokine production and complement regulation, independent of external complement sources or stressors. We show that decreased *CFH* levels and activity result in increased levels of inflammatory cytokines, chemokines and growth factors, that are known to play a role in AMD, as well as several other neurodegenerative diseases. Although our study reported here does not delineate between the two protein products made by the *CFH* gene (*i*.*e*. FH and FHL-1) it is reasonable to assume that given FH is the major splice variant expressed by hTERT-RPE1 cells (see Fig. S1) that the biological consequences of *CFH* gene silencing in our study are mediated primarily by FH.

Based on the levels of secreted inflammatory proteins and PCA analysis (Fig. 3), we observed a clear segregation between control RPE cells (siNeg) and RPE cells deprived of *CFH*. The main discriminatory factors were IL-6 and IL-8, which were also the most upregulated cytokines after *CFH* silencing. Besides their role in inflammation, IL-6 and IL-8 are both members of the senescence-associated secretory pathway (SASP) and involved in ageing processes. Indeed, H2O2-mediated senescence in ARPE19 cells leads to increased levels of IL-6 and IL-8 when FH levels were reduced (39). Moreover, increased systemic IL-6 levels were found in patients with AMD, mostly in relation to the late subtypes of the disease (40). Interestingly, a study exploring potential new drug targets for AMD identified IL-6 as a candidate target (41).

We found in RPE cells lacking FH increased secreted levels of GM-CSF (Fig. 3), a growth factor that promotes activation and survival of microglia cells and macrophages (42). Interestingly, GM-CSF levels have been found to be elevated in the vitreous of postmortem human eyes genotyped for the *CFH* Y402H SNP, and in parallel accumulation of choroidal macrophages was observed (43). In this study, local accumulation of GM-CSF was found after stimulation with the anaphylatoxins C3a and C5a (43). Our data suggest that RPE cells may be a source of GM-CSF found in the *CFH* Y402H post-mortem eye.

Serpin E1, also known as Plasminogen Activator Inhibitor-1 (PAI-1), was upregulated in siCFH RPE cells. Serpin E1 is involved in ECM remodeling and angiogenesis (44), processes that are altered at the Bruch’
ss membrane/choroid interface in AMD (2). Serpin E1 is also considered a senescence marker in several tissues (44). High levels of Serpin E1 have been associated with neovascularization in AMD and diabetic retinopathy (45). Serpin E1 mediates some of its effects *via* binding to the α5β3 integrin (46), and interestingly also FH and its truncated form FHL-1 modulate RPE function *via* binding a closely related integrin, α5β1 (47). Other factors altered by FH in RPE cells include CXCL1/GROα, a chemokine responsible for neutrophil recruitment and activation (48) that has been found increased in aqueous humor of AMD patients (49), and MIP-1α and MIP-1β, which have been found to be involved in inflammation-mediated damage in the retina (50). FH loss in RPE cells also leads to upregulation of IL-1ra, which has been found to be highly expressed by RPE cells in response to IL-1β and TNFα stimulation (51). Interestingly, IL-1β has been found to be highly expressed in iPSC-derived RPE cells carrying the *CFH* 402H variant (52) and TNFα accumulates in the BrM and choroid in eyes from *CFH* 402H donors (53).

Our results are in line with independent observations that in AMD, as well as in the presence of the *CFH* 402H variant, inflammation is increased. However, the signaling pathways involved in the regulation of inflammation in RPE cells are not fully characterized, and most importantly the pathway by which FH regulates inflammation in RPE cells was not known. The majority of cytokines differentially regulated in FH-deprived conditions are target of the NF-κB pathway. NF-κB plays a central role in regulating cellular responses to inflammation and is often activated in concert with the complement system. NF-κB consists of transcription factor complexes expressed in most cell types and can be activated in response to a variety of stimuli or stressors, which allow the cell to respond and adapt to variations in the microenvironment including infections, growth factor levels or oxidative stress (32). The most common target genes of the NF-κB pathway are inflammatory cytokines, responsible for recruitment of neutrophils and macrophages at the inflamed site (38): cells that are also recruited after local complement activation (17). We have shown here, that NF-κB activation follows suppression of *CFH* expression, which in turn results in an upregulation of NF-κB dependent cytokines. Reducing NF-κB levels leads to a reduction in the expression and secretion of most of the upregulated cytokines, including IL-6, IL-8, CCL2, Serpin E1, GM-CSF and CXCL1/GROα. Consequently, the cytokine profile of the siCFH + siRELA group clusters tightly with the siNeg control.

The interaction between the complement system and the NF-κB pathway has been reported in other cell types and pathologies, mostly in the context of cell response to complement-mediated damage. Here, NF-κB is suggested to play a pro-survival role. Mouse fibroblasts, HELA cells and HEK293 cells lacking NF-κB p65, are all more sensitive to complement-mediated damage. Here, the NF-κB pathway was found to suppress JNK-dependent programmed necrotic cell death, rather than being involved in complement regulation and inflammation, since no changes in the expression or activity of the complement regulators CD46, CD55 and CD59 were observed (54). Also, HUVEC cells and human coronary endothelial cells (ECs) show NF-κB activation in response to MAC formation, in a AKT/endosomes dependent mechanism (55). Data from *in vitro* and *in vivo* models of Alzheimer disease (AD) support the hypothesis that astroglia, rather than neurons, are the principle site of NF-κB overactivation and those cells are then primarily responsible for the NF-κB -dependent increase in C3. Importantly in this study, NF-κB binding sites were confirmed in the *C3* promoter. Moreover, the NF-κB /C3 axis in astroglia cells was suggested to be dependent on the classical complement, rather than alternative complement pathway, due to changes in gene expression of C1q and C4 and not Cfb and Cfh (56). Interestingly, HIV infection activates NF-κB in astrocytes and promotes C3 production in a IL6-dependent manner (57). Also, results from kidney *in vivo* models of renal injury, provide evidence that the NF-κB pathway plays an important role in renal damage mediated by enhanced local complement activation (58).

It is important to note that RPE cells, in contrast to all these previously mentioned cell types, show a significant level of tolerance to complement-mediated damage (59). This may explain why they do not exploit the NF-κB pathway to respond to external complement stimulation, but rather regulate this pathway to maintain physiological levels of complement and inflammatory factors. Here, the NF-κB pathway was not seen activated as pro-survival pathway to respond to a complement activation mediated insult, since addition of neither C3 nor C3b had an impact on NF-κB activity. However, previous studies have reported that RPE cells show NF-κB activation in response to oxidative stress. For instance, ARPE19 cells show increased phosphorylation in NF-κB p65 in response to either short or long exposure to H2O2 (60). In our previous study we show that FH loss increases oxidative and metabolic stress, both stressors of which may induce NF-κB activation as a survival mechanism. We have also shown that genes involved in oxidative stress response and mitochondrial stability (*e*.*g*. PGC1α) were differentially regulated by complement activation: interestingly these are targets of the NF-κB pathway as well. Further studies will be necessary to understand the mechanism by which FH loss leads to NF-κB activation. Being co-expressed in RPE cells, FH could directly modulate NF-κB pathway activation on the protein level. Alternatively, the absence of FH as an antioxidant factor could activate the NF-κB pathway in the context of oxidative stress response in RPE cells.

The observation, that a subset of cytokines (MIP-1α, MIP-1β and IL-1ra), which were increased upon FH loss, were not reduced after silencing of NF-κB p65 suggests, that additional pathways are likely involved in the interplay between complement and NF-κB pathway. Several pathways have been described as being involved in the homeostasis of RPE cells, which could be regulated by FH. For example, knock-out of CXCR5 in RPE cells leads to an AMD-like phenotype and transcriptome profile highlights the role of PI3K-Akt and mTOR signaling, as important pathways for RPE homeostasis (61). Another possibility involves the regulation of the transcription factor AP1, which has been found to be regulated together with NF-κB in response to blue-light mediated damage (62).

Although the exact mechanism remains to be discovered, our data contribute to the understanding around how risk alleles in *CFH* which result in reduced FH/FHL-1 activity may increase the risk for AMD. We suggest that in RPE cells, FH and the NF-κB pathway work in synergy to maintain cellular homeostasis, keeping both pro-inflammatory pathways in balance and check (summarized in Fig. 5). In case either one of them is dysregulated due to genetic risk, age and/or local stressors, the RPE microenvironment changes towards a pro-inflammatory AMD-like phenotype, with NF-κB as well as the alternative complement pathway acting as major protagonists.

**Figure 5.**
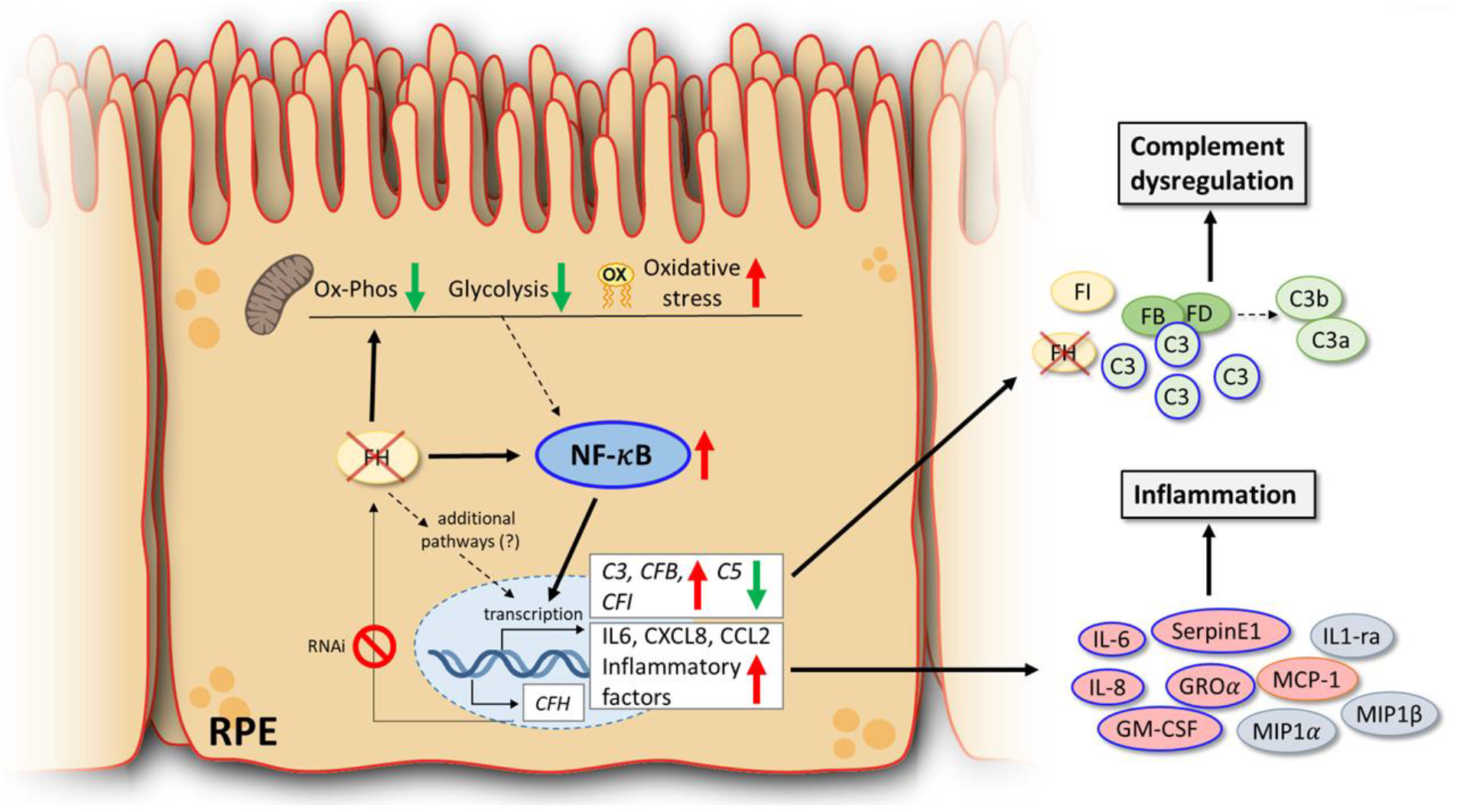
Summary schematic of NF-kB-mediated inflammation driven by the loss of *CFH*. Reduced levels of *CFH* via RNAi in RPE cells leads to activation of the NF-κB pathway (blue). The NF-κB pathway regulates gene transcription of inflammatory cytokines (light red) as well as positive (green) and negative (yellow) regulators of complement activation, secreted from RPE. FH-deprived RPE cells are characterized by reduced levels of oxidative phosphorylation (Ox-Phos) and glycolysis, as well as increased oxidative stress and oxidized lipids levels (ox). Metabolic and oxidative stresses are known positive regulators of the NF-κB pathway and could contribute to the activation of the NF-κB pathway in FH-reduced conditions (dotted arrow). Secreted proteins circled in blue are directly modulated through the actions of the NFkB pathway. Cytokines secreted in FH-deprived RPE cells, which are not regulated by NF-κB pathway, are labeled grey.

